# Time- and performance-dependent enhancement of tactospatial working memory caused by online anodal tDCS of the posterior parietal cortex

**DOI:** 10.64898/2026.06.08.730789

**Authors:** Miro Grundei, Luca Kämmer, Timo Torsten Schmidt, Till Nierhaus, Felix Blankenburg

## Abstract

**Background:** Spatial working memory (WM) relies on posterior parietal cortex (PPC) within a distributed fronto-parietal network, yet whether online anodal transcranial direct current stimulation (tDCS) of PPC modulates tactile WM, and how behavioral effects unfold over time, remains unclear.

**Objective:** We tested whether online anodal tDCS over left PPC modulates performance in a tactospatial WM task and whether stimulation effects vary over time and as a function of baseline performance.

**Methods:** In a double-blind, sham-controlled, within-subject crossover design, 32 healthy adults completed two counterbalanced sessions (Active, Sham). Each session comprised baseline, stimulation, and post-stimulation phases while participants performed a retro-cued delayed match-to-sample task with vibrotactile patterns delivered to the left index finger. During the stimulation phase, participants received either 2 mA anodal or sham tDCS over left PPC for 15 min, depending on session. Behavioral accuracy was analyzed using a sliding-window approach combined with cluster-based permutation testing.

**Results:** Active stimulation induced a significant, gradual improvement in WM accuracy relative to sham, emerging approximately eight minutes after stimulation onset and corresponding to a ∼5% performance increase (Hedges’ g = 0.43). Stimulation effects were baseline-dependent, with individuals showing lower initial WM performance exhibiting larger behavioral gains, whereas high performers showed minimal benefits.

**Conclusions:** These findings show that online anodal stimulation of left PPC can modulate tactile WM performance in a time- and baseline-dependent manner. More broadly, the results suggest that behavioral effects of parietal tDCS emerge gradually during ongoing stimulation and depend on initial cognitive state.

**Highlights:**

- Online anodal PPC-tDCS enhances tactile working memory
- Behavioral gains emerge after ∼8 min of stimulation
- Active stimulation improves accuracy by ∼5% versus sham
- Lower baseline performers show larger stimulation benefits
- Sliding-window analyses capture dynamic online tDCS effects

## Introduction

Working memory (WM) refers to the capacity-limited system that maintains and manipulates information over brief periods, supporting essential cognitive functions such as reasoning, planning, and decision-making (Baddeley, 1974; 2012). Classical and contemporary models of WM describe a distributed fronto-parietal network that supports the maintenance and flexible prioritization of behaviorally relevant information across sensory modalities, with posterior parietal regions often implicated in representing spatial and sensory-specific properties and frontal regions in more abstract, task-dependent control processes (e.g., Curtis & D’Esposito, 2003; D’Esposito & Postle, 2015; Christophel et al., 2017; Schmidt & Blankenburg, 2018). Although WM processes generalize across modalities, most studies investigating the behavioral effects of non-invasive brain stimulation on WM have focused on the visual domain.

Non-invasive brain stimulation techniques such as transcranial direct current stimulation (tDCS) have been widely used to probe causal contributions of specific cortical regions to cognition. In anodal tDCS, the positive electrode is placed over the region of interest, slightly depolarizing cortical tissue and increasing neuronal excitability (Nitsche & Paulus, 2000), which can modulate behavioral performance in WM tasks (Fregni et al., 2005; Zaehle et al., 2011; Tseng et al., 2021). However, behavioral effects of tDCS on WM are often heterogeneous across studies and appear to depend on stimulation target, task demands, stimulation timing, and inter-individual differences in baseline cognitive state (Hsu et al., 2014; Heinen et al., 2016; Tseng et al., 2021; Vergallito et al., 2022). In particular, several studies suggest that individuals with lower baseline performance often exhibit larger behavioral gains in response to tDCS (Hsu et al., 2014; Heinen et al., 2016; Tseng et al., 2021; Vergallito et al., 2022).

Consistent with its proposed role in representing sensory-specific and spatial WM content, evidence from both neuroimaging and brain-stimulation studies suggests that posterior parietal cortex (PPC) is a particularly promising target for modulating *tactospatial* WM performance. In the tactile domain, pioneering work in non-human primates demonstrated robust WM-related activity during vibrotactile delayed match-to-sample paradigms (Romo & Salinas, 2001; Romo et al., 2012), while human studies provided evidence for the maintenance and updating of vibrotactile WM representations within distributed fronto-parietal networks (Spitzer et al., 2010; Spitzer & Blankenburg, 2011; Spitzer & Blankenburg, 2012; Schmidt et al., 2017; Uluç et al., 2018; Wu et al., 2018). Recent neuroimaging work has shown that PPC preferentially represents tactospatial WM information during maintenance (Schmidt & Blankenburg, 2018; Schmidt et al., 2021; Grundei et al., 2025), and that left PPC activity predicts behavioral performance during tactile WM (Grundei et al., 2025). Consistent with these observations, causal studies using single-pulse transcranial magnetic stimulation (TMS) have shown that transient disruption of PPC impairs tactile and cross-modal WM performance, supporting a causal role of parietal cortex in somatosensory memory processing (Ku et al., 2015a; Ku et al., 2015b). These findings complement related perturbation studies implicating somatosensory and prefrontal cortices in tactile WM maintenance (Zhao et al., 2018). Beyond transient interference approaches, brain stimulation studies in the visual domain further suggest that excitatory parietal stimulation may enhance WM performance more reliably than prefrontal stimulation in some task contexts (Wang et al., 2019), although recent findings also highlight substantial variability in behavioral outcomes (Jiang et al., 2024). However, it remains unknown whether online excitatory stimulation of PPC can dynamically enhance tactile WM performance over the course of ongoing task execution.

In addition to the targeted brain region, the temporal dynamics of ongoing stimulation may critically influence behavioral effects. Physiological studies have shown that anodal tDCS effects accumulate gradually over the course of stimulation, with robust excitability changes emerging only after several minutes of continuous current application (Nitsche & Paulus, 2001; Nitsche & Paulus, 2011). However, relatively few studies have examined how behavioral effects evolve during ongoing stimulation, particularly in non-visual WM paradigms.

In this double-blind, sham-controlled, within-subject crossover study, we investigated whether anodal tDCS over the left PPC modulates performance in a well-controlled tactile delayed match-to-sample task (Schmidt & Blankenburg, 2018; Grundei et al., 2025). Participants completed two counterbalanced sessions (Active and Sham tDCS), each comprising baseline, stimulation, and post-stimulation phases. To capture temporally evolving stimulation effects during ongoing task performance, we applied a sliding-window analysis combined with cluster-based permutation testing. Based on previous evidence linking PPC activity to tactile WM performance, the gradual accumulation of anodal tDCS effects, and baseline-dependent variability in stimulation responsiveness, we hypothesized that anodal stimulation of left PPC would produce a time-dependent enhancement of tactile WM performance during stimulation, with larger behavioral benefits in participants showing lower baseline performance.

## Materials and Methods

### Participants

Thirty-two healthy adults (19 female, 12 male, 1 diverse; mean age = 25.97 ± 3.36 years, range: 21–36) participated in the study. Sample size was determined based on previous within-subject tDCS studies investigating WM, which commonly included approximately 20– 25 participants (e.g., Tseng et al., 2012; Heinen et al., 2016; Wang et al., 2019), with N = 32 chosen to provide comparable or greater statistical sensitivity for the present design. All participants reported normal or corrected-to-normal vision and no history of neurological or psychiatric conditions. Additional exclusion criteria included prior epileptic seizures, migraine, psychoactive medication, recent alcohol consumption, recent night-shift work, or use of substances affecting cognition within 24 hours. All participants provided written informed consent, and the study was approved by the ethics committee of Freie Universität Berlin (013/2020).

### Tactile Working Memory Task

To probe tactospatial WM, participants completed a vibrotactile delayed match-to-sample task based on Schmidt & Blankenburg (2018) with a short WM delay (Grundei et al., 2025). Two successive vibrotactile stimuli were delivered to the left index finger (700 ms each), followed by a masking stimulus (500 ms) and a visual retro-cue (display ‘1’ or ‘2’) indicating which of the stimuli to retain in memory during a 7 s delay phase. Stimuli were presented via a 16-pin (4×4) piezoelectric Braille display (QuaeroSys, Germany; Figure 1A) where each pin vibrated with a 30 Hz sinusoidal amplitude modulation, resulting in the formation of smooth and surface-like spatial patterns with equalized overall energy. The two sample stimuli were selected to be maximally dissimilar (Pearson’s correlation r < 0.05) and the full-field masking stimulus followed to eliminate somatosensory perceptual residues. At retrieval, participants performed a two-alternative forced-choice discrimination between the remembered target and a *foil* pattern moderately similar to the target (r = 0.6–0.7). At each trial, stimulus patterns were drawn at random from 200 possible stimulus sets each consisting of two sample stimuli and a corresponding foil with the respective correlational properties. Participants responded via right hand key-presses within a 1500 ms response window and received feedback (500 ms display ‘+’, ‘–’, or non-response ‘×’). Inter-trial intervals were jittered between 2–4 s and each run consisted of 24 trials, corresponding to ∼6.5 min per run.

**Figure 1.**
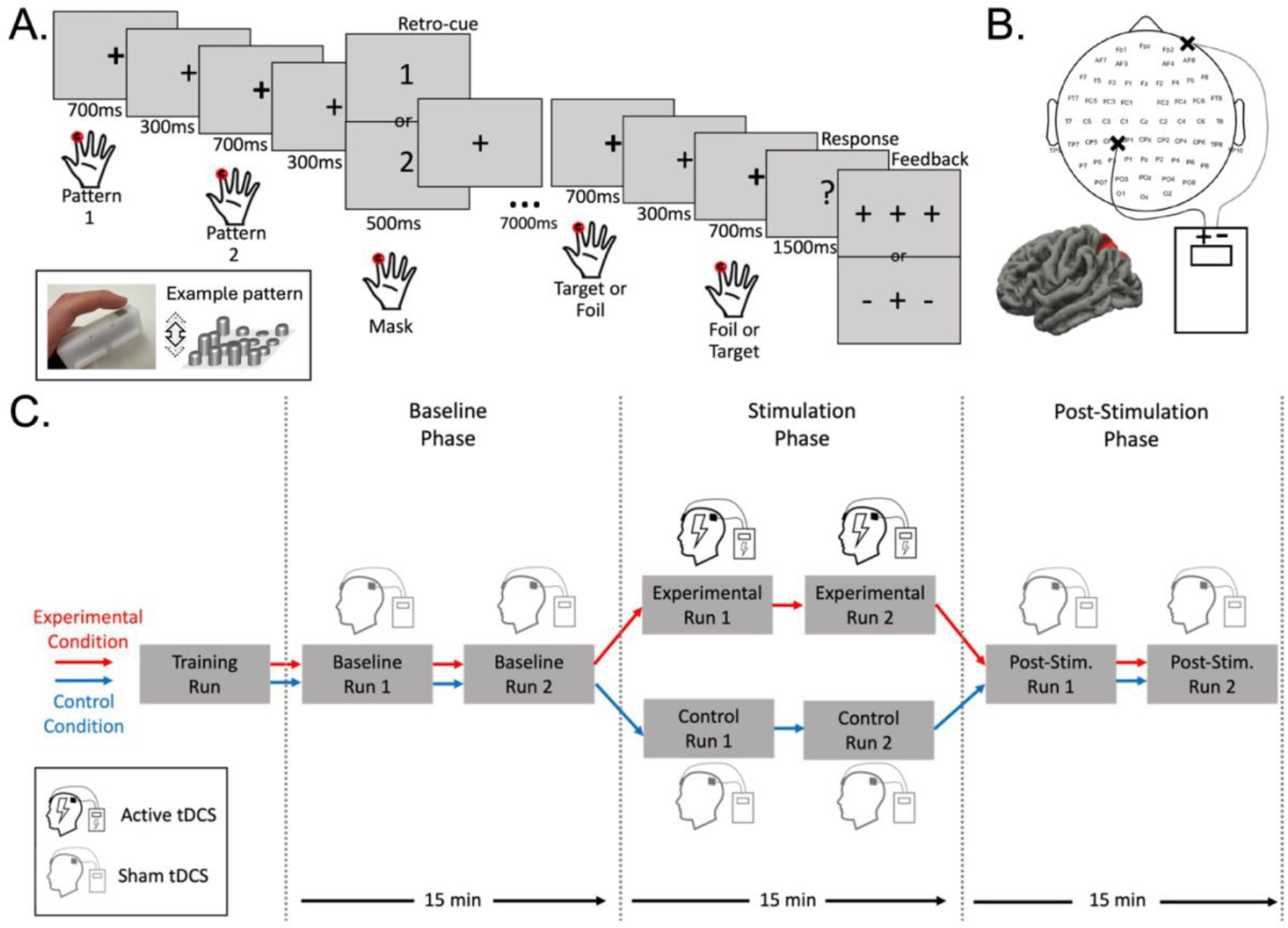
Experimental Paradigm and Procedure. A) Retro-cued delayed match-to-sample task. Each trial started with a presentation of two consecutive vibrotactile sample stimuli followed by a mask stimulus (vibration of the full stimulus display at maximum amplitude). A retro-cue (‘1’ or ‘2’) was presented together with the mask to indicate which of the two sample stimuli had to be retained for the 7 s WM delay period. After the delay phase participants were presented with two stimuli, one of which was the memorized stimulus (target) and the other one was a changed version of it (foil). Participants indicated with a right-hand key-press which of the two stimuli was the target. Small rectangular panel on the left: Vibrotactile Braille-like pin display presented to the left index finger. Example stimulus pattern presented using a 4 x 4 pin matrix on which pins vibrated with different amplitudes but with the same frequency (30 Hz). Amplitudes were modelled to be smooth and surface-like. Figure adapted from Schmidt and Blankenburg (2018) and Grundei et al. (2025). B) The stimulation sites where the electrodes were fixed to the scalp. The anode was fixed between the 10–20 coordinates CP3 and CP1. Below that is a depiction of a brain with the area (left PPC) that was targeted with the anodal tDCS highlighted in red. C) Stimulation procedure of the experiment. The general setup of the experiment is the same in both testing sessions. After familiarization with the paradigm through a training run, the participants completed two baseline runs, two runs in the stimulation phase, and two post-stimulation runs. During the baseline phase the participants received sham stimulation while a baseline of their performance was established. In the stimulation phase the participants received either sham or active tDCS stimulation, depending on the session. In the post-stimulation phase the participants again received sham stimulation. Each phase consisting of two runs took 15 minutes.

### Experimental Procedure and tDCS Protocol

All participants completed both an active and a sham session on separate days, with the order counterbalanced. Each session consisted of: (1) a practice run, (2) two baseline runs (sham stimulation), (3) two stimulation-phase runs (active or sham tDCS), and (4) two post-stimulation runs (sham). In the sham session, all runs used sham stimulation. The experimental procedure is depicted in Figure 1C. The study followed a double-blind protocol in which both participants and experimenters were blinded using device-generated codes that encrypted the programmed stimulation sequence.

Transcranial direct current stimulation (tDCS) was delivered using a neuroConn DC-Stimulator (neuroCare Group GmbH, Munich, Germany). A 4.3 cm diameter rubber anode (18.49 cm2) was placed over the left PPC, localized between positions CP1 and CP3 of the 10–20 Electroencephalography (EEG) cap (Figure 1B), which project onto the left posterior parietal cortex at approximately Talairach coordinates X = −34, Y = −48, Z = 58. Left PPC was selected based on previous neuroimaging findings using the same tactile WM paradigm, demonstrating performance-relevant stimulus representations in this region (Schmidt & Blankenburg, 2018; Grundei et al., 2025). The cathode was positioned at the forehead over the right supraorbital area.

### Integrity of the Blind

Following completion of both experimental sessions, participants were verbally debriefed regarding the stimulation procedure and reported being unable to distinguish between the different sessions, indicating successful blinding.

### Experimental Design and Statistical Analysis

Analyses were performed in Python using NumPy, pandas, SciPy, and Matplotlib. No participants or trials were excluded.

Accounting for the fact that behavioral effects of online tDCS likely evolve gradually during ongoing stimulation rather than align with predefined run boundaries, accuracy was computed using a sliding-window approach rather than run-wise averaging which collapses arbitrary intervals and provides only coarse temporal resolution. The sliding window contained 24 trials, corresponding to the number of trials in a run. This window size balances two considerations: (i) providing sufficiently many trials for a stable estimate of accuracy, given the binary nature of the measurement, and (ii) preserving temporal resolution to detect stimulation-locked changes across runs in stimulation and post-stimulation phases of the experiment. For each participant and condition (Sham and Active), sliding-window averaging produced a smoothed time series of accuracy estimates, with one sample per window shift. These time series naturally exhibit temporal autocorrelation because adjacent windows overlap. Importantly, the subsequent cluster-based permutation test fully accounts for this dependence structure: the same sliding-window operation is applied to every permuted dataset, ensuring that the empirical null distribution preserves the autocorrelation inherent in the raw data. Thus, statistical inference remains valid despite non-independence of adjacent timepoints, consistent with procedures widely used in time-resolved M/EEG analysis (Maris & Oostenveld, 2007).

To test for stimulation-related changes, we computed within-subject difference time series (Active−Sham). Statistical testing focused on the a priori defined stimulation and post-stimulation intervals, restricting inference to those temporal windows in which stimulation effects could potentially be expected. For each timepoint in this interval, a one-sample t-statistic (equivalent to a paired t-test) was computed across participants. Timepoints exceeding a one-sided cluster-forming threshold of p < 0.05 (uncorrected) were grouped into temporally contiguous clusters. Cluster-level significance was assessed using a sign-flip permutation procedure (10,000 iterations). For each permutation, the sign of each participant’s entire difference time series was randomly inverted, implementing the null hypothesis that Active and Sham labels are exchangeable within subjects. Because sign-flipping preserves the entire temporal structure of each participant’s trajectory, temporal autocorrelation is explicitly retained in the null distribution, preventing inflated false-positive rates. For each permuted dataset, the largest cluster mass (sum of t-values) was recorded, forming a family-wise–error-controlled distribution. Observed clusters were assigned corrected p-values by comparing their masses against this distribution, with clusters considered significant at p < 0.05. For descriptive purposes, effect sizes (mean difference, 95% CI, Cohen’s d, and Hedges’ g) were computed using the non-permuted data, whereas all inference was based exclusively on the cluster-level permutation statistics. Hedges’ g was computed as the primary effect size as it provides an inherent correction for inflation by small sample sizes (Hedges & Olkin, 2014).

Individual differences in baseline cognitive ability have been shown to moderate the behavioral effects of non-invasive brain stimulation. Several tDCS studies have reported that individuals with lower initial WM performance tend to exhibit larger stimulation-related benefits, whereas high-performing individuals often show little or no improvement (e.g., Hsu et al., 2014; Gözenman & Berryhill, 2016; Heinen et al., 2016; Tseng et al., 2021; Vergallito et al., 2024). Based on previous reports of baseline-dependent responsiveness to tDCS, we assessed whether baseline task performance modulated the stimulation effect observed in the cluster-based permutation analysis. Baseline performance was defined for each participant using the pre-stimulation portion of the task (averaged across runs 1 and 2 of both sessions). To quantify each participant’s stimulation response, we used the cluster-level summary statistic derived from the within-subject cluster-based permutation analysis. Baseline-dependence was first assessed continuously by correlating baseline accuracy with the stimulation effect (Active– Sham cluster-level summary statistic) using Pearson correlation. For visualization purposes, participants were additionally assigned to “low-baseline” and “high-baseline” groups using a median split, and stimulation effects were compared between groups using one-sided Welch’s t-test, consistent with the a priori directional hypothesis that lower-baseline participants would exhibit larger stimulation benefits.

## Results

### Tactile Working Memory Performance

Participants successfully performed the demanding tactile WM task, as indicated by an overall accuracy of 62.32 ± 6.34% (mean ± SD; session 1: 62.07 ± 7.07%; session 2: 62.56 ± 6.90%). To assess potential learning effects prior to sorting trials into Active and Sham stimulation conditions, we conducted a 2 × 6 repeated-measures ANOVA with the factors session (1, 2) and run (1–6). We found no systematic performance differences between sessions (main effect of session: F(1,31) = 0.14, p = 0.71) or across runs (main effect of run: F(5,155) = 0.81, p = 0.55). The session × run interaction was also not significant (F(5,155) = 1.11, p = 0.36), indicating that performance remained stable across sessions and runs. Additionally, controlling for potential presentation-order effects, repeated-measures ANOVAs showed that performance was not significantly influenced by whether the first or second sample stimulus was memorized (Session 1: F(1,31) = 0.76, p = 0.39; Session 2: F(1,31) = 0.46, p = 0.50).

### Modulation of Tactile Working Memory Performance by tDCS

The sliding-window analysis revealed how behavioral performance evolved over time during active and sham stimulation (Figure 2A). During the stimulation phase, the Active-tDCS condition showed a distinct upward shift relative to Sham, emerging approximately eight minutes after stimulation onset. The cluster-based permutation test revealed a significant positive cluster indicating higher accuracy during Active stimulation (p = 0.043; 5.3% improvement in accuracy; Cohen’s d = 0.44; Hedges’ g = 0.43). This effect was robust to reasonable variation in the sliding-window length, remaining significant across window sizes ranging from 21 to 38 trials, indicating that the finding was not contingent on a specific smoothing parameter.

**Figure 2.**
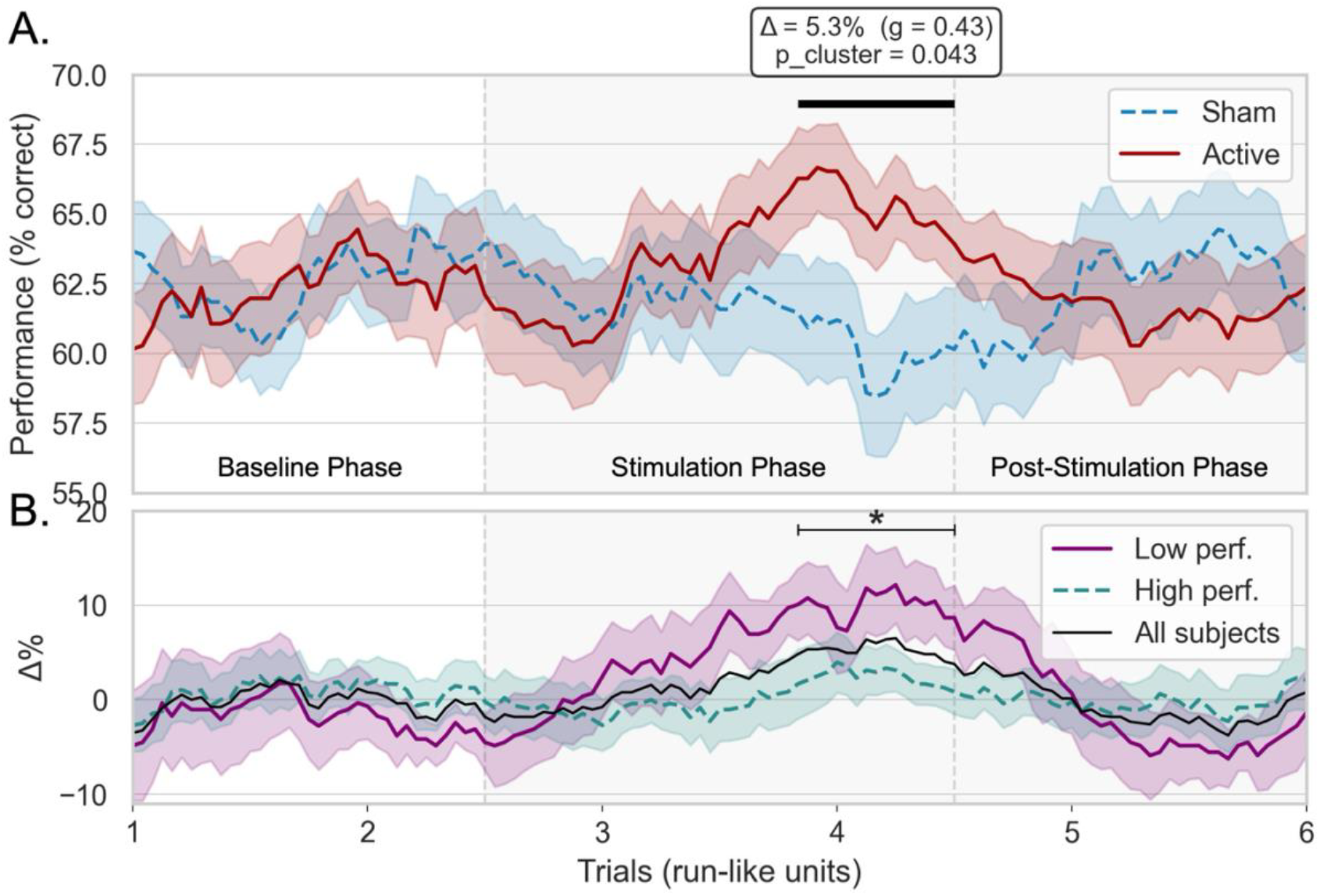
Running mean of participants’ performance using a sliding window approach. Shadings correspond to standard error of the mean. Window size: 24 trials. A) Performance for Active (red line) and Sham (blue line, dashed) stimulation conditions during baseline phase (left, spanning runs 1 and 2), stimulation phase (middle, grey shading, spanning runs 3 and 4) and post-stimulation phase (right, grey shading, spanning runs 5 and 6). Thick black line on top indicates significant time-points of the cluster-based permutation test (p < 0.05). B) Performance time-series of the Active–Sham difference (Δ%) of all subjects (black line) as well as participants assigned to the Low (purple line) and High (teal line, dashed) baseline groups for visualization of the baseline-dependent stimulation effect. Asterisk indicates significant t-contrast (p < 0.05) between performer groups within the significant Active tDCS effect.

### Baseline-Dependent Variability in Stimulation Effects

Because behavioral responses to tDCS commonly vary as a function of individual baseline performance, we next examined whether the observed stimulation benefit scaled with baseline task ability. Consistent with our hypothesis, baseline performance negatively correlated with the magnitude of the stimulation effect when treated as a continuous variable (r = −0.33, p = 0.034), indicating that lower baseline performance was associated with larger behavioral gains during active stimulation. For visualization purposes, participants were additionally separated into low- and high-baseline groups using a median split (Figure 2B). Consistent with the continuous analysis, participants with lower baseline performance exhibited a significantly larger stimulation benefit than participants with higher baseline performance (t(27.3) = 1.75, p = 0.048), reflected in a pronounced positive Active–Sham difference during the significant stimulation interval.

## Discussion

The present study investigated whether anodal tDCS over the left PPC modulates performance in a demanding tactile WM task using a double-blind, sham-controlled, within-subject crossover design. Participants completed a delayed match-to-sample paradigm in which vibratory spatial patterns delivered to the left index finger had to be retained over a brief delay. Each of the two sessions consisted of six runs, structured into baseline, stimulation, and post-stimulation phases. During the stimulation phase, participants received either active anodal or sham tDCS over the left PPC. Using a time-resolved sliding-window approach combined with cluster-based permutation testing, we identified a significant stimulation-related enhancement of behavioral performance that emerged approximately eight minutes after stimulation onset. Importantly, stimulation effects were baseline-dependent, with individuals showing lower tactile WM performance at baseline exhibiting larger behavioral gains during active stimulation.

The present study was designed to characterize how behavioral effects of online tDCS evolve over the course of ongoing stimulation, with the baseline phase providing a measure of initial task performance and the post-stimulation phase allowing assessment of potential aftereffects. Conducting active and sham sessions on separate days prevented carryover effects, and implementing a double-blind protocol minimized expectancy-related biases. While standard run-wise ANOVAs can fail to detect temporally dynamic stimulation-induced changes because tDCS acts as a continuously evolving neuromodulatory process, our sliding-window analysis provides a time-resolved metric which is more sensitive to non-stationary behavioral changes. The cluster-based permutation approach used here preserves temporal autocorrelation and controls family-wise error, providing a robust statistical method for identifying stimulation-locked effects that would remain obscured in run-averaged analyses.

The delayed onset of the observed behavioral effect aligns well with physiological work showing that tDCS-induced excitability changes accumulate gradually, reaching robust strength only after several minutes of ongoing stimulation (Nitsche & Paulus, 2001; Nitsche & Paulus, 2011). Although the observed effect size (g = 0.43) corresponds to a small-to-medium enhancement (Cohen, 2013), it is notably larger than the average effect of prefrontal tDCS on WM reported in recent meta-analytic syntheses, which indicate only small improvements (g ≈ 0.14; Wischnewski et al., 2024). The use of a double-blind crossover design consistent with current best practices for non-invasive brain stimulation studies (Bikson et al., 2018) lends additional support to the conclusion that the behavioral gains represent a small but genuine tDCS effect. The ∼5% accuracy improvement observed here is in line with behavioral gains found in several visual WM studies applying anodal tDCS (e.g., Fregni et al., 2005; Berryhill & Jones, 2012; Jones & Berryhill, 2012). However, most previous work has focused on prefrontal stimulation in the visual modality, where meta-analyses show inconsistent and often modality-specific outcomes, with more reliable effects for verbal than visuospatial WM (Mancuso et al., 2016; Wischnewski et al., 2024). Building on previous TMS studies demonstrating that transient disruption of PPC impairs tactile and cross-modal WM performance (Ku et al., 2015a; Ku et al., 2015b), the present findings show that online anodal PPC-tDCS can conversely enhance tactile WM performance during ongoing task execution.

The baseline-dependence of stimulation efficacy observed in the present study is consistent with a growing body of evidence indicating that behavioral responses to tDCS depend strongly on individual factors, including cortical state, neuroanatomy, and especially initial task performance, where lower-performing individuals often exhibit larger behavioral gains during stimulation (Krause & Cohen Kadosh, 2014; Vergallito et al., 2022). Specifically, our analyses were motivated by a study which investigated the impact of 20 minutes of anodal active versus sham tDCS over PPC on visual WM performance (Tseng et al., 2012). While the authors did not detect a main effect of stimulation, after categorizing participants into low- and high-performer groups based on the median score from the sham condition, performance gains were identified for the low-performance group. Consistent with this account, in the current study we similarly find that individuals with low baseline performance show higher WM improvements in response to PPC stimulation during a tactile WM task.

The present findings are consistent with a functional contribution of left PPC to tactile WM performance and converge with previous neuroimaging work using the same delayed match-to-sample paradigm showing that tactospatial WM content can be decoded from parietal activation patterns (Schmidt & Blankenburg, 2018; Grundei et al., 2025). Notably, Grundei et al. (2025) demonstrated a temporal progression in representational dynamics: stimulus information first emerged in somatosensory cortex and was subsequently expressed in PPC, with only this later PPC decoding correlating with behavioral accuracy. Together with a decoding study showing that left PPC flexibly recodes tactile information into task-dependent, categorical WM representations (Velenosi et al., 2020), these results highlight the functional relevance of PPC for successful tactile WM and underscore its flexible role in maintaining and transforming somatosensory information. Beyond content representation, PPC interacts with premotor regions to support task-dependent processing: premotor cortex drives the rehearsal of tactospatial WM content in parietal cortex, whereas vibrotactile frequency information is maintained via prefrontal–premotor pathways (Schmidt et al., 2021). These findings from the tactile modality parallel evidence from visual WM, where decoding and lesion studies demonstrate that PPC supports the maintenance and transformation of spatial mnemonic content: Stimulus-specific information such as orientation, location, shape and motion can be reliably decoded during delay periods (Christophel et al., 2017), PPC encodes increasingly abstract and behaviorally relevant representations (Christophel et al., 2017; Chunharas et al., 2024; Rademaker et al., 2019) and parietal damage selectively impairs the precision of spatial WM (Mackey et al., 2016). Overall, our causal findings integrate with this broader evidence to establish the PPC as a key node supporting the maintenance and transformation of WM representations across multiple modalities.

Interestingly, the observed behavioral enhancement was restricted to the stimulation phase and did not persist into the post-stimulation interval. While this temporal profile differs from studies reporting prolonged aftereffects of anodal stimulation in motor cortex (Nitsche & Paulus, 2001), behavioral effects in higher-order association cortices may depend more strongly on ongoing task engagement and network state. In this context, the present findings suggest that online modulation of task-relevant parietal networks may be sufficient to transiently enhance behavioral performance, even in the absence of measurable post-stimulation effects.

Despite our encouraging findings, several limitations should be noted. First, the present design does not fully dissociate WM-specific effects from more domain-general changes in cognitive state, such as arousal or sustained attention, as stimulation was applied continuously during task performance and no independent vigilance control task was included. Although the delayed onset of the behavioral effect and the anatomically motivated targeting of left PPC argue against a purely nonspecific account, future studies incorporating active control tasks or independent measures of vigilance will be important for clarifying the specificity of the observed effects. Second, overall task accuracy was relatively low (∼63%), likely attributable to the high similarity between target and foil patterns (60–70% correlation). Future studies may benefit from adjusting task difficulty or using adaptive procedures to maintain performance at more a balanced range (≈70–75%) for detecting enhancement effects. Moreover, binary accuracy measures introduce substantial trial-wise variability and incorporating continuous or graded responses could provide more sensitive indices of stimulation-related changes. Third, although electrode placement targeted left PPC using standardized EEG coordinates, individual differences in skull thickness, gyrification, and tissue conductivity likely introduced variability in the delivered electric fields, a known limitation of conventional tDCS (Datta et al., 2009; Laakso et al., 2015; Thielscher et al., 2026). MRI-guided neuronavigation or individualized E-field modeling would improve stimulation focality and interpretability (Meinzer et al., 2024). Finally, combining non-invasive brain stimulation with fMRI or MEG may help elucidate how stimulation-induced changes in cortical excitability influence fronto-parietal WM networks.

In summary, the present findings demonstrate that online anodal stimulation of the left PPC can enhance tactospatial WM performance. The observed behavioral gains emerged gradually, depended on baseline ability, and converged with neurophysiological evidence implicating PPC in the maintenance and transformation of somatosensory information. Together, these results extend prior WM-tDCS research into the tactile domain and demonstrate that online anodal stimulation of left PPC can modulate tactile WM performance in a time- and baseline-dependent manner.

## Acknowledgements

This research was funded by the German Research Foundation (project grants to FB: Research Unit 5429/1 (467143400), BL 977/4-1).

